# A linear discriminant analysis model of imbalanced associative learning in the mushroom body compartment

**DOI:** 10.1101/2022.09.23.508775

**Authors:** David Lipshutz, Aneesh Kashalikar, Shiva Farashahi, Dmitri B. Chklovskii

## Abstract

To adapt to their environments, animals learn associations between sensory stimuli and unconditioned stimuli. In invertebrates, olfactory associative learning primarily occurs in the mushroom body, which is segregated into separate compartments. Within each compartment, Kenyon cells (KCs) encoding sparse odor representations project onto mushroom body output neurons (MBONs) whose outputs guide behavior. Associated with each compartment is a dopamine neuron (DAN) that modulates plasticity of the KC-MBON synapses within the compartment. Interestingly, DAN-induced plasticity of the KC-MBON synapse is imbalanced in the sense that it only weakens the synapse and is temporally sparse. We propose a normative mechanistic model of the MBON as a linear discriminant analysis (LDA) classifier that predicts the presence of an unconditioned stimulus (class identity) given a KC odor representation (feature vector). Starting from a principled LDA objective function and under the assumption of temporally sparse DAN activity, we derive an online algorithm which maps onto the mushroom body compartment. Our model accounts for the imbalanced learning at the KC-MBON synapse and makes testable predictions that provide clear contrasts with existing models.

## 1 Introduction

Behavioral responses of animals are shaped in part by learned associations between sensory stimuli (e.g., odors) and unconditioned stimuli (e.g., sugar, water or heat). An important challenge in neuroscience is to understand the neural mechanisms that underlie associative learning. In invertebrates, the mushroom body is a well studied brain region that plays a central role in olfactory associative learning [1, 2, 3, 4]. The goal of this work is to propose a normative, mechanistic model of associative learning in the mushroom body that accounts for experimental observations and provide clear contrasts with existing models.

The mushroom body is segregated into functionally independent compartments [5], Figure 1. Within each compartment, Kenyon cells (KCs), which encode sparse odor representations [6], form synapses with the dendrites of mushroom body output neurons (MBONs), whose outputs guide learned behavior [7]. Associated with each compartment is a single Dopamine neuron (DAN) that responds to an unconditioned stimuli [8, 9], and projects its axon into the mushroom body compartment where it innervates the KC-MBON synapses to modulate plasticity, implicating the KC-MBON synapse as the synaptic substrate for associative learning in invertebrates.

**Figure 1:**
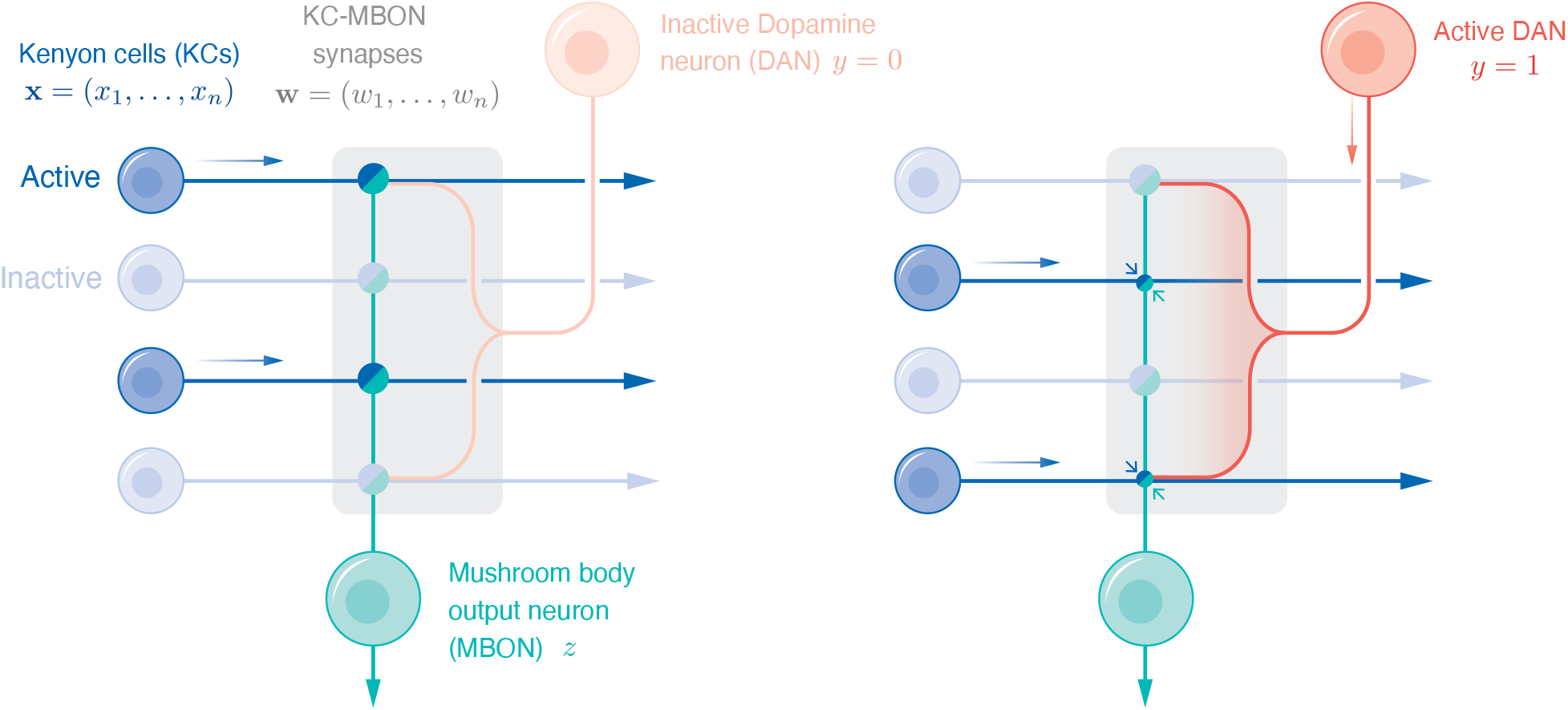
A simplified schematics of a mushroom body compartment illustrating that co-activation of the KCs and DAN weakens the KC-MBON synapse. In both the left and right panels, the mushroom body compartment, indicated by the shaded box, is innervated by the axons of multiple KCs, the dendrites from one MBON and the axon terminals of one DAN. The bi-colored circles at the intersections of the KC axons and the MBON dendrites denote the KC-MBON synapses. Faintly shaded cell bodies indicate inactive neurons and boldly shaded cell bodies indicate active neurons. In the left panel, the DAN is inactive. In the right panel, the DAN is active and co-activation of the KCs and the DAN weakens the associated KC-MBON synapses (as illustrated by the smaller synapses).

Experimental evidence suggests that learning at the KC-MBON synapse is imbalanced in the sense that DAN-induced plasticity is one-sided and temporally sparse. In particular, co-activation of a KC and the DAN *weakens* the KC-MBON synapse (see Figure 1, right) and DAN-induced plasticity is independent of the MBON activity [5]. This suggests that DAN-induced plasticity is one-sided and some other mechanism such as homeostatic plasticity is responsible for strengthening the KC-MBON synapse. Furthermore, since each DAN responds to one type of unconditioned stimulus [2], which only constitutes a small fraction of all unconditioned stimuli, the DAN activity is temporally sparse.

In this work, we propose a normative, mechanistic model of associative learning in the mushroom body that accounts for the imbalanced learning. We model each MBON as a linear discriminant analysis (LDA) classifier, which predicts if an associated unconditioned stimulus is present (the class label) given a KC odor representation (the feature vector). Under this interpretation, the KC-MBON synapses and an MBON bias term define a hyperplane in the high-dimensional space of KC odor representations that separates odor representations associated with the unconditioned stimulus from all other odor representations, Figure 2.

**Figure 2:**
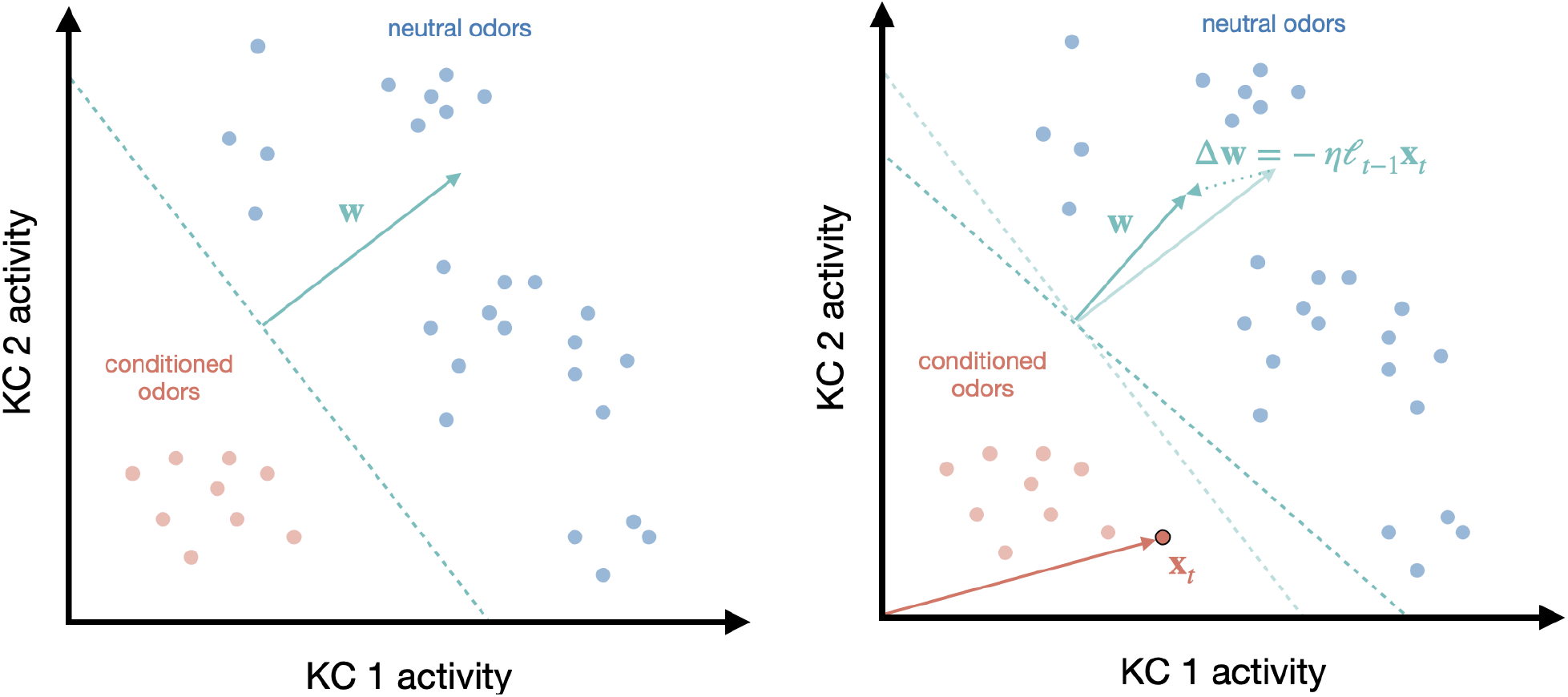
Left: Illustration of the hyperplane ℋ = {**x** : **w**^*⊤*^**x** = *b*} (dashed teal line) in the space of KC activities that separates KC responses to conditioned odors from KC responses to neutral odors. Each light red (resp. blue) dot denotes the KC response to a conditioned (resp. neutral) odor; that is, the set of vectors **x**_*t*_ such that *y*_*t*_ = 1 (resp. *y*_*t*_ = 0). The teal arrow denotes the vector of KC-MBON synaptic weights **w**, which is translated to show that it is orthogonal to the hyperplane ℋ. Right: Illustration of the weakening of the synaptic weights **w** in response to co-activation of the KCs and the DAN. The KC activities **x**_*t*_ are denoted by the dark red dot with black border. The change of the synaptic weights ∆**w** is in the direction −**x**_*t*_ and is proportional to *ℓ*_*t−*1_, the time elapsed since the last presentation of a conditioned odor. The hyperplane ℋ rotates to remain orthogonal to **w**. Note that the change in bias ∆*b*, which translates the hyperplane, is not depicted in this illustration.

Here, ‘normative’ refers to the fact that our mechanistic model is interpretable as an algorithm for optimizing an LDA objective. The normative approach is top-down in the sense that first the circuit objective is proposed and then an optimization algorithm is derived and compared with known physiology. There are several advantages to this approach. First, it directly relates the circuit objective to its mechanism; for example, neural activities and synaptic weight updates are interpretable as steps in an algorithm for solving a relevant circuit objective. Second, the approach distills down what aspects of the physiological are essential for optimizing the circuit objective and what aspects are not captured by the objective. Third, normative models are often analytically tractable, which allows them to be analyzed for any input statistics without resorting to exhaustive numerical simulation.

To derive our algorithm, we start with a convex objective for LDA (in terms of the KC-MBON synaptic weights). The objective can be optimized in the offline setting by taking gradient descent steps with respect to the KC-MBON synaptic weights. To obtain an online algorithm that accounts for the imbalanced learning, we take advantage of the fact that DAN activity is temporally sparse to obtain online approximations of the input statistics. Finally, we show numerically that our algorithm performs well even when DAN activity is not temporally sparse.

Our model makes testable predictions that are a direct result of the learning imbalance. First, our model predicts that DAN-induced plasticity at the KC-MBON synapse is sensitive to the time elapsed since the DAN was last active. Second, our model predicts that if the DAN is never active, then the KC-MBON synapses adapt to align with the mean KC activity (normalized by the covariance of the KC activity).

## 2 Results

### Problem statement

Consider a simplified mushroom body compartment that consists of KC axons, the axon terminals from one DAN and the dendrites of one MBON, Figure 1. Let *n* denote the number of KCs and **x**_1_, …, **x**_*T*_ ∈ ℝ^*n*^ be a sequence of *n*-dimensional vectors whose components represent the activities of the KCs. Let *y*_1_, …, *y*_*T*_ ∈ {0, 1} be a sequence indicating if the DAN is active or inactive at a given time. We assume the DAN is active if and only if the unconditioned stimulus is present. If the DAN is active at time *t* (i.e., *y*_*t*_ = 1), we refer to **x**_*t*_ as a *conditioned odor* (representation), whereas if the DAN is inactive at time *t* (i.e., *y*_*t*_ = 0), we refer to **x**_*t*_ as a *neutral odor* (representation).

We consider the case that the MBON is a linear classifier that predicts the class label *y*_*t*_ at time *t* given the KC activities **x**_*t*_. Let **w** be an *n*-dimensional vector whose components represent the strength of the KC-MBON synapses. We assume that for each *t*, the activity of the MBON is given by

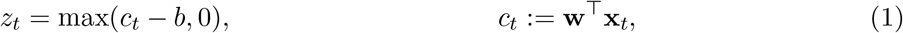

where *b* represents the ‘bias’ of the MBON. Under this interpretation, the KC-MBON synapses **w** and MBON bias *b* define a hyperplane ℋ := {**x** : **w**^*⊤*^**x** = *b*} in the *n*-dimensional space of KC activities that separates conditioned odors from neutral odors, Figure 2. In this case, *z*_*t*_ *>* 0 (resp. *z*_*t*_ = 0) corresponds to the prediction *y*_*t*_ = 0 (resp. *y*_*t*_ = 1). In other words, the MBON is active when predicting there is no unconditioned stimulus and inactive when predicting there is an unconditioned stimulus, which is consistent with experimental observations [2].

To be consistent with the experimental observations in [5], we require that co-activation of a KC and the DAN weakens the KC-MBON synapse. Mathematically, this can be represented as follows:

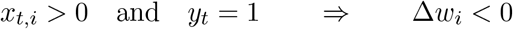

Furthermore, we require that DAN-induced plasticity at the KC-MBON synapse is independent of the MBON output *z*; that is, if *y* = 1, then ∆**w** does not depend on *z*. Finally, we require that the DAN activity is sparse in time. Letting *π*_0_ and *π*_1_ denote the proportion of *t* such that *y*_*t*_ = 0 and *y*_*t*_ = 1, respectively, this is mathematically equivalent to the following:

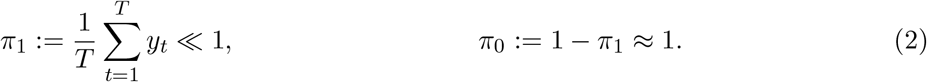

We seek a normative online algorithm for learning the KC-MBON synaptic weights **w** and bias *b* which is consistent with physiological observations.

### Linear discriminant analysis

In this work, we model the mushroom body compartment as an LDA classifier. LDA is a linear statistical method for classification [10, section 4.3], which makes the following simplifying assumption: the conditional probability distributions *p*(**x**|*y* = 0) and *p*(**x**|*y* = 1) are both Gaussian with common *n* × *n* covariance matrix **Σ**; that is

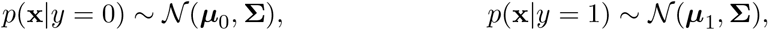

where ***µ***_0_ and ***µ***_1_ denote the means of the class 0 and class 1 feature vectors. In this case, the optimal decision criteria for assigning class 0 (resp. class 1) to feature vector **x** is **w**^*⊤*^**x** *> b* (resp. **w**^*⊤*^**x** *< b*), where

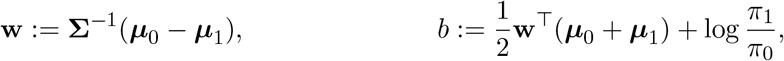

and *π*_*i*_ denotes the probability that a samples belongs to class *i*, for *i* = 0, 1. In particular, the hyperplane ℋ = {**x** ∈ ℝ^*n*^ : **w**^*⊤*^**x** = *b*} defines the optimal separation boundary for predicting whether a feature vector belongs to class 0 or class 1. While LDA assumes a specific generative model, it performs well in practice even when the assumptions do not hold [11].

There are a number of existing online LDA algorithms [12, 13, 14], and LDA is equivalent to canonical correlation analysis [15] for which there are online algorithms with biologically plausible neural network implementations [16, 17, 18]. However, these algorithms do not naturally map onto the mushroom body compartment. In the next section we derive an online LDA algorithm that maps onto the mushroom body compartment and matches experimental observations.

### An LDA objective function and offline algorithm

We can write the LDA problem as the solution of the following convex minimization problem:

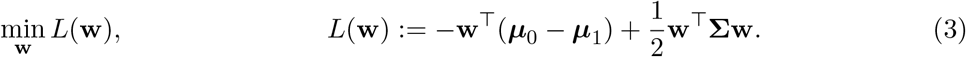

Differentiating *L*(**w**) with respect to **w** we see that the unique minimum is given by

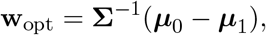

where we are assuming that **Σ** is full rank.

In the offline setting, we can optimize the strongly convex objective (3) by running the gradient descent algorithm:

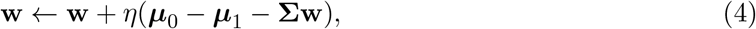

where *η >* 0 is the step size. Formally, taking the step size *η* to zero yields the linear gradient flow

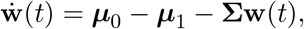

whose solution is given by

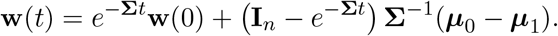

In particular, we see that the solution **w**(*t*) converges exponentially to the optimal solution **w**_opt_. In the next section, we use the fact that the unconditioned stimuli are sparse in time to derive an online algorithm that will map onto the mushroom body circuit.

### An online algorithm for imbalanced learning

In the online setting, the class means ***µ***_0_, ***µ***_1_ and the covariance **Σ** are not available. Instead, at each time *t* the algorithm has access to the feature vector **x**_*t*_ and class label *y*_*t*_. To derive our online algorithm, we make online approximations of the offline quantities ***µ***_0_, ***µ***_1_ and **Σ** that are based on the fact that the unconditioned stimuli are sparse in time, see equation (2). First, we note that we can rewrite the sample class means

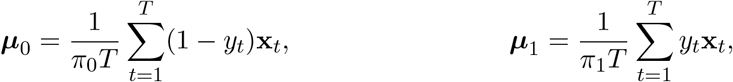

and the sample covariance

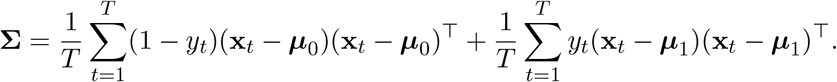

### Estimating the mean response to a neutral odor and the covariance

Since *π*_0_ ≈ 1, we can keep a running approximation of ***µ***_0_ by performing the following update

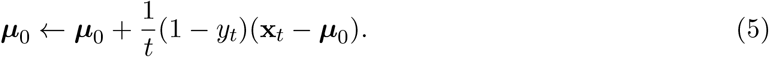

In addition, since *y*_*t*_ = 0 for ‘most’ samples, we can replace **Σ** with the online approximation

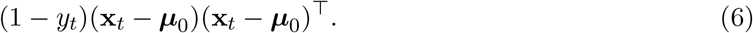

Setting *ζ* := **w**^*⊤*^***µ***_0_ and multiplying the update in equation (5) on the left by **w**^*⊤*^, we can keep a running approximation of *ζ* by performing the following update

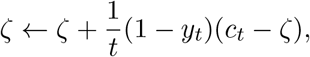

where we recall that *c*_*t*_ := **w**^*⊤*^**x**_*t*_. Right multiplying the online approximation of **Σ** in equation (6) by **w** and using the definitions for *c*_*t*_ and *ζ*, we can replace **Σw** with the online approximation

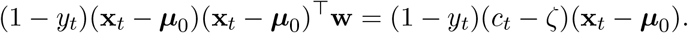

### Estimating the mean response to a conditioned odor

To obtain an online approximation of ***µ***_1_, we first note that 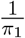 is approximately equal to the average time elapsed between class 1 samples. To see this, let *t*_1_, …, *t*_*r*_ denote the subset of times such that *y*_*t*_ = 1. Then, letting *t*_0_ = 0, we have

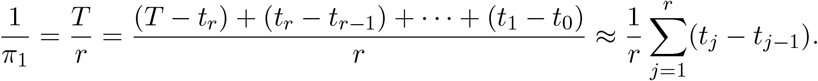

Thus, in the online setting, when the *j*^th^ class 1 sample is presented (i.e., *y*_*t*_ = 1), we can use the time elapsed since the last class 1 sample, *t*_*j*_ − *t*_*j−*1_, as an online estimate of 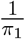. Setting *ℓ*_0_ = 1 and

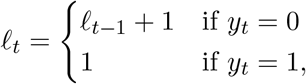

we see that at time *t* such that *y*_*t*_ = 1, *ℓ*_*t−*1_ denotes the time elapsed since the last class 1 sample, so 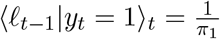. Assuming that the variables *ℓ*_*t*_ and **x**_*t*_ are independent given *y*_*t*_ = 1 — i.e., the KC representation **x**_*t*_ of a conditioned odor is independent of the time elapsed since the last conditioned odor *ℓ*_*t−*1_ — we see that ***µ***_1_ = ⟨*ℓ*_*t−*1_|*y*_*t*_ = 1⟩_*t*_⟨**x**_*t*_|*y*_*t*_ = 1⟩_*t*_ = ⟨*ℓ*_*t−*1_**x**_*t*_|*y*_*t*_ = 1⟩_*t*_. Thus, given *y*_*t*_ = 1, we can replace ***µ***_1_ with the online approximation *ℓ*_*t−*1_**x**_*t*_.

### Estimating the bias

To estimate the bias *b*, we note that because *π*_0_ ≈ 1 and 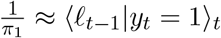

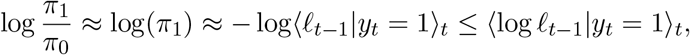

where the final inequality follows from the fact that log is concave and Jensen’s inequality (with equality holding when the variance of *ℓ*_*t−*1_ given *y*_*t*_ = 1 is zero). Thus, assuming the variance of the time elapsed between conditioned odors is small, we can estimate the bias *b* in the online setting with the updates:

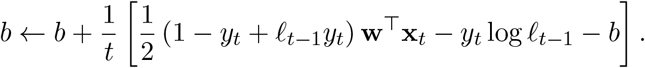

Substituting these approximations into the offline update rules in equation (4) yields our online algorithm (Algorithm 1).

#### Algorithm 1: LDA in the mushroom body compartment

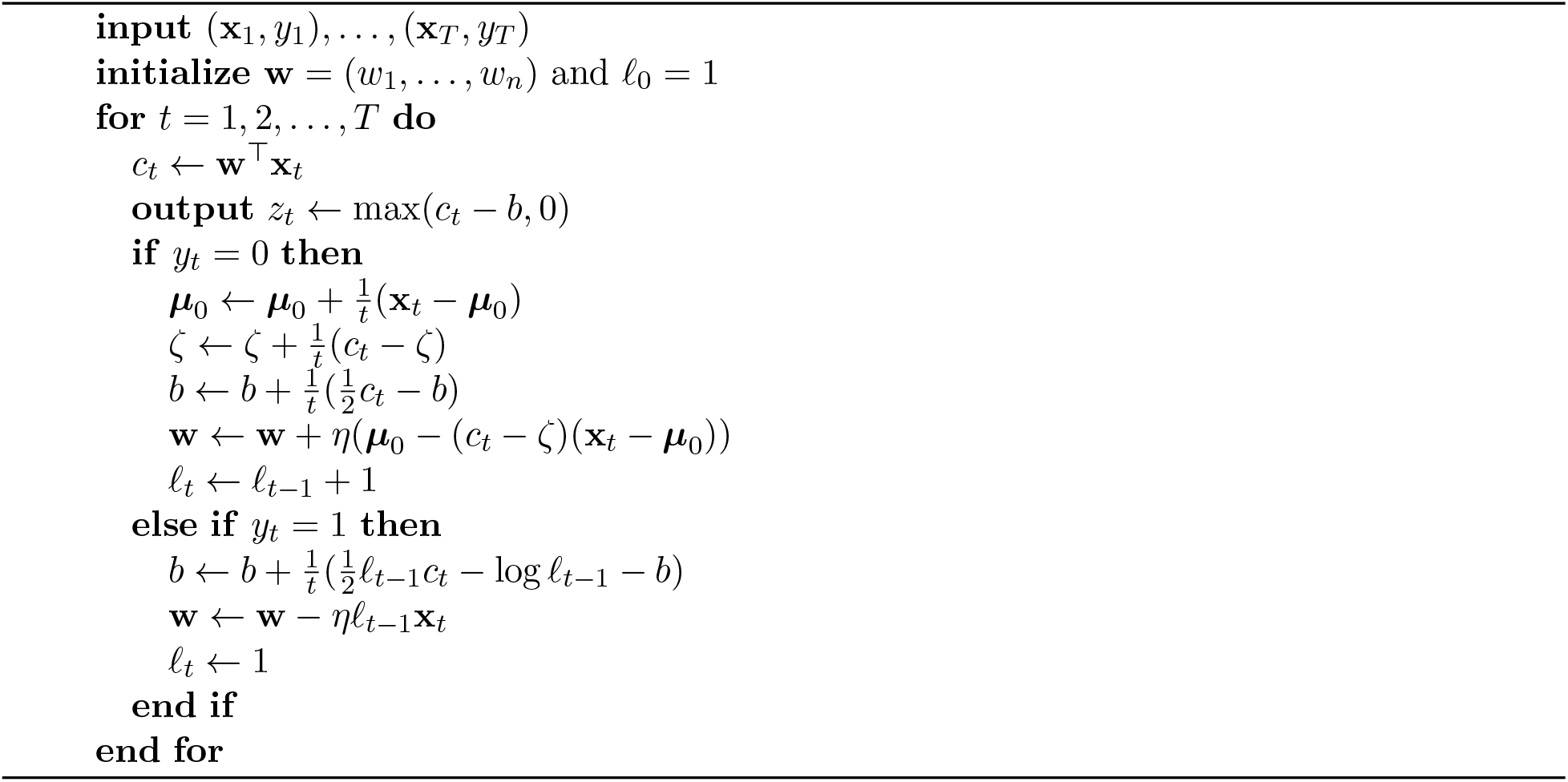

In view of Jensen’s inequality, if the variance of the time elapsed between conditioned odors is large, then the bias *b* will be overestimated, meaning that the MBON will be less active than optimal. In other words, irregular intervals between DAN activity biases the MBON to be less active (i.e., predict that the unconditioned stimulus is present more often).

### Relation to the mushroom body

#### Biological interpretations

Algorithm 1 naturally maps onto the mushroom body compartment. In this case, *w*_*i*_ represents the synaptic weight between KC *i* and the MBON, for *i* = 1, …, *n*, and *b* represents the ‘bias’ of the MBON, which denotes the threshold of the soma. The scalar *c*_*t*_ can be represented as dendritic current. The running sample mean of the KC activities can be stored as calcium concentrations in the pre-synaptic KC-MBON terminals and the projected mean *ζ* can be stored as calcium concentration in the post-synaptic KC-MBON terminals. The scalar *ℓ*_*t−*1_ can be represented as the sensitivity of the KC-MBON synapses to DAN-induced plasticity, which is proportional to the time elapsed since the DAN was last active.

Hige et al. [5] showed that learning each compartment is hetergeneous in the sense the compartments have separate timescales for learning. This heterogeneity can be accounted for in our model by using a different the learning rate *η >* 0 for each compartment, which interestingly is the only hyper-parameter in our model.

#### Model predictions

In our model, when the DAN is active (i.e., *y*_*t*_ = 1), the synapses are updated as follows:

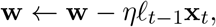

which is inline with experimental evidence showing that DAN-induced plasticity is independent of the MBON activity *z* and co-activation of the KCs and the DAN reduces the strength of the synapses between the KCs and the MBON [5]; see Figure 2 (right) for a geometric interpretation of the plasticity rule. Moreover, the update is proportional to *ℓ*_*t−*1_, which results in our first prediction.

*The strength of DAN induced plasticity in the KC-MBON synapses is proportional to the time elapsed since the last time the DAN was active*.

When the DAN is not active (i.e., *y*_*t*_ = 0), the synapses are updated according to the homeostatis rule

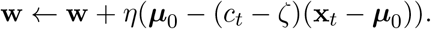

In particular, when the DAN is not active, the average KC-MBON synaptic weight update is **w** ← **w** + *η*(***µ***_0_ − **Σ**_0_**w**), where we have used the fact that ⟨(*c*_*t*_ − *ζ*)(**x**_*t*_ − ***µ***_0_)⟩_*t*_ = **Σ**_0_**w**. This yields our second prediction.

*In the absence of DAN activity, the KC-MBON synaptic weights will align with the mean KC activity (normalized by the covariance of the KC odor representations); that is*, **w** *will converge to* 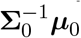.

Our 2 predictions are precisely the reason our algorithm can learn using plasticity rules for the KC-MBON synapses that are independent of the MBON activity, which is consistent with experimental observations [5] and in contrast to existing models [19, 20, 21].

### Numerical experiments

In this section, we test Algorithm 1 on synthetic and real datasets. We test our algorithm on inputs when our assumption *π*_1_ ≪ 1 holds, but also on inputs when *π*_1_ ≈ 0.5. To evaluate our algorithm, we measure the running accuracy of the projections *z*_*t*_ over the previous min(1000, *t*) iterations, where the algorithm is accurate at the *t*^th^ iterate if *z*_*t*_ = 0 and *y*_*t*_ = 1 or if *z*_*t*_ *>* 0 and *y*_*t*_ = 0.

#### Synthetic dataset

We begin by evaluating Algorithm 1 on a synthetic dataset generated by a mixture of 2 overlapping Gaussian distributions, so that the optimal accuracy is less than 1. The data points of the 2 classes are each drawn from a 2-dimensional mean with common covariance. We simulate datasets of 10^5^ data points using the same mean and covariance in both classes but vary the frequency of class 1 samples encountered. We consider the cases *π*_1_ = 0.1, 0.2, 0.3, 0.4, 0.5. In Figure 3 (left) we plot the error and the accuracy of our model for varying *π*_1_. Remarkably, while the derivation of Algorithm 1 relied on the fact that *π*_1_ ≪ 1, the algorithm still performs well even when *π*_1_ = 0.5

**Figure 3:**
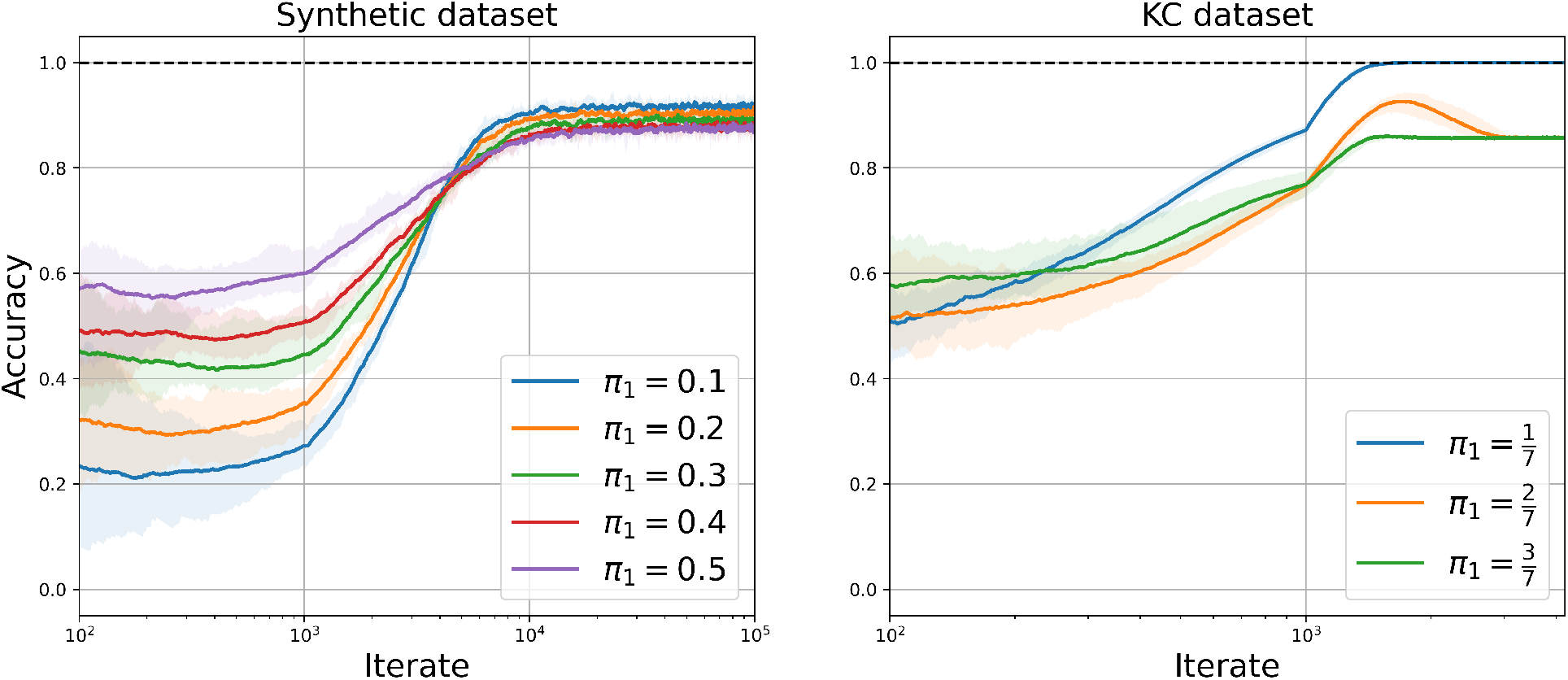
Performance of Algorithm 1 on the synthetic datasets (left) and the KC dataset (right). Each line denotes the mean accuracy over 10 runs. Each shaded region indicates the area between the minimum and maximum accuracy over 10 runs.

#### KC activities dataset

We test our model on KC activities reported in [22]. They recorded odor-evoked KC responses in the fly mushroom body. The dataset we tested on contains the responses of 124 KCs in a single fly to the presentation of 7 odors, see [22, Figure 1]. To ensure the KC responses are well conditioned, we add Gaussian noise with covariance *ϵ***I**_124_, where *ϵ* = 0.01. We apply Algorithm 1 to the KC dataset. We first consider the case that odor 1 denotes the class 1 odor and odors 2–7 denote the class 0 odors, so 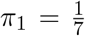. We then consider the cases that odors 1–2 (resp. 1–3) odors denote the class 1 odors and the remaining odors denote the class 0 odors, so 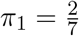 (resp. 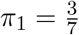). In Figure 3 (right) we plot the error and accuracy of our model for varying *π*_1_. Impressively, the modified algorithm performs well (approximately 85% accuracy) even when the assumption *π*_1_ ≪ 1 is violated.

#### Competing MBONs

Using the KC activities dataset, we model 2 MBONs with competing valences by running 2 instances of Algorithm 1 in parallel with different class assignments for the odors. We consider the case that odor 1 is aversive, odor 7 is attractive and the remaining odors are neutral. For MBON 1 (resp. MBON 2), we assume that odor 1 (resp. odor 7) denotes the class 1 odor and odors 2–7 (resp. odors 1–6) denote the class 0 odors, so that MBON 1 (resp. MBON 2) activity promotes approach (resp. avoid) behavior. Let *z*_*i,t*_ denote the output of MBON *i* ∈ {1, 2}. At each iterate *t*, if odor 1 (resp. odor 7, odors 2–6) is presented, then the model is accurate if *z*_1,*t*_ *>* 0 and *z*_2,*t*_ = 0 (resp. *z*_1,*t*_ = 0 and *z*_2,*t*_ *>* 0, resp. *z*_1,*t*_ *>* 0 and *z*_2,*t*_ *>* 0), and inaccurate otherwise. We then repeated the experiment two more times, but with odor 2 (resp. odor 3) labeled as aversive and odor 6 (resp. odor 5) as attractive. In Figure 4, we plot the performance of the competing MBONs.

**Figure 4:**
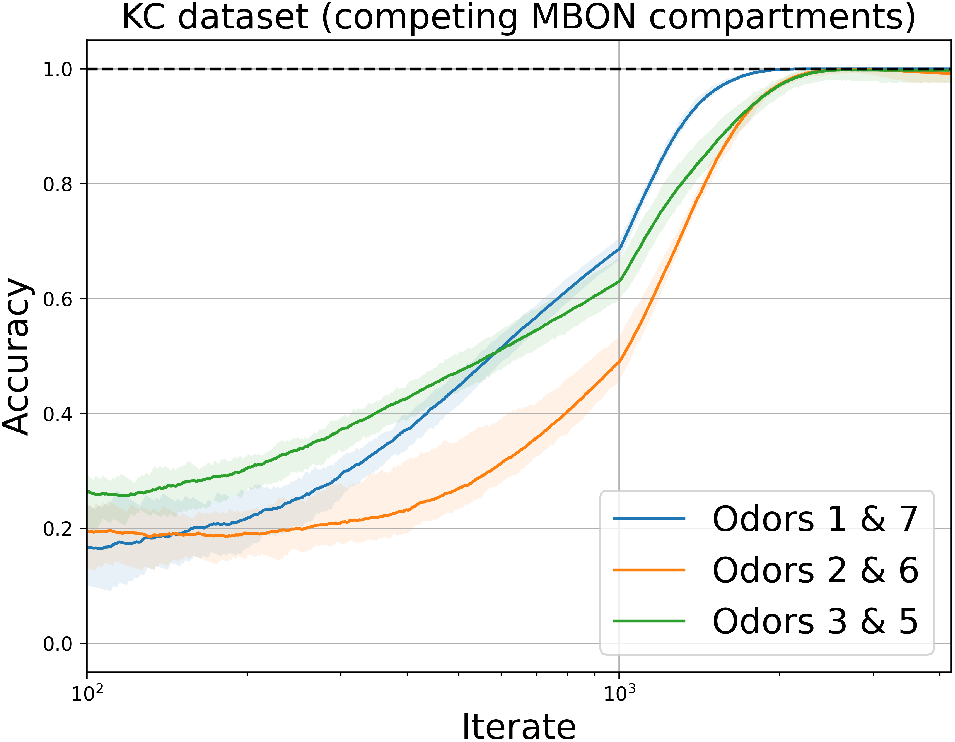
Performance of parallel runs of Algorithm 1 on the KC dataset with different class assignments for each run. Each line denotes the mean accuracy over 10 runs. Each shaded region indicates the area between the minimum and maximum accuracy over 10 runs.

## 3 Methods

### Details of numerical experiments

The experiments were performed on an Apple iMac with a 3.2 GHz 8-Core Intel Xeon W processor. For each experiment, we used a learning rate of the form

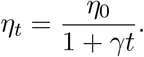

We chose the parameters *η*_0_ and *γ* by performing a grid search over *η*_0_ ∈ {1, 10^*−*1^, 10^*−*2^, 10^*−*3^, 10^*−*4^} and *γ* ∈ {10^*−*2^, 10^*−*3^, 10^*−*4^, 10^*−*5^, 10^*−*6^}. The optimal parameters for the synthetic dataset (resp. KC dataset) are *η*_0_ = 10^*−*1^ and *γ* = 10^*−*3^ (resp. *η*_0_ = 10^*−*1^ and *γ* = 10^*−*4^).

## 4 Discussion

### Summary

In this work, we proposed a normative, analytically tractable LDA model of the mushroom body compartment that accounts for imbalanced learning at the KC-MBON synapse. Our model makes testable predictions that provide clear contrasts with existing models of associative learning in the mushroom body. Interestingly, our model of plasticity and the KC-MBON synapse does not depend on the MBON activity, but rather predicts that DAN-induced plasticity is sensitive to the time elapsed since the last time the DAN was active. Finally, testing our model on synthetic and real datasets shows that it performs well under a variety of conditions.

### Relation to existing models

There are a number of existing computational models of associative learning in the mushroom body, many of which are faithful to biophysical details and successfully capture important computational principles underlying associative learning in the mushroom body. Huerta and Nowotny [19], Bazhenov et al. [20], Huerta et al. [23], Smith et al. [24], Peng and Chittka [25] propose computational models of associative learning in the mushroom body that capture known physiologically details (see, e.g., [19, p. 3]) and accurately learn associations between sensory inputs and unconditioned stimuli. Through extensive numerical simulations, their computational models can explain a number of phenomena. For example, Huerta and Nowotny [19] show that the organization of the mushroom body supports fast and robust associative learning, Bazhenov et al. [20] show that interactions between unsupervised and supervised forms of learning can explain how the timescale of associative learning depends on experimental conditions, and Peng and Chittka [25] show how complex forms of learning (e.g., peak shift) depend on different mechanistic aspects of learning in the mushroom body. In this work, we propose a top-down normative model of learning at the KC-MBON synapse, which contrasts with the bottom-up approach in these works that build models closely tied to physiological evidence. In this way, the our model is interpretable as an algorithm for optimizing a circuit objective and the output can be predicted analytically for any environmental condition without needing to resort to numerical simulation. In addition, our normative model makes testable predictions that are in clear contrast with these models, providing a method for validation or invalidation of our model.

In addition to these models, Bennett et al. [21] propose a reinforcement learning model of the KC-MBON synapses as minimizing reinforcement prediction errors. They first consider a model in which the reinforcement signal is computed as the difference between DAN activities, so their plasticity rule requires 2 DANs to innervate a single mushroom body compartment, which is in contrast to experiment evidence showing that most compartments only receive inputs from a single DAN [9]. To account for this experimental observation, they propose a heuristic modification that adds a constant source of synaptic potentiation, which can be viewed as a form of homeostatic plasticity and is inline with experimental evidence. However, the modification is not normative and can fail to minimize prediction errors. One of the main challenges they encounter is how to balance homeostatic plasticity with DAN-induced plasticity. Our model provides a normative account for balancing homeostatic plasticity and DAN-induced plasticity: DAN-induced plasticity depends on the time-elapsed since the DAN was last active.

### Limitations

Our model is a dramatic simplification of the mushroom body focused on providing a normative account of learning at the KC-MBON synapse that can account for how balance between DAN-induced plasticity and homeostatic plasticity is maintained. Consequently, our does not account for a number of the physiological details that are captured by existing models. Furthermore, there are addition features such as feedback connections in the mushroom body that have been recently discovered and are relevant for associative learning [7, 26], which are also not capture by our model.

## Acknowledgements

We are grateful to Lucy Reading-Ikkanda for creating Figure 1. We thank Yanis Bahroun, Siavash Golkar, Jason Moore and Tiberiu Tesileanu for helpful feedback on an earlier draft of this work.

## Notes

### Competing Interest Statement

The authors have declared no competing interest.

## References

[1] Martin Heisenberg. Mushroom body memoir: from maps to models. Nature Reviews Neuroscience, 4(4):266–275, 2003.

[2] David Owald and Scott Waddell. Olfactory learning skews mushroom body output pathways to steer behavioral choice in Drosophila. Current Opinion in Neurobiology, 35:178–184, 2015.

[3] Katharina Eichler, Feng Li, Ashok Litwin-Kumar, Youngser Park, Ingrid Andrade, Casey M Schneider-Mizell, Timo Saumweber, Annina Huser, Claire Eschbach, Bertram Gerber, et al. The complete connectome of a learning and memory centre in an insect brain. Nature, 548 (7666):175, 2017.

[4] Mehrab N Modi, Yichun Shuai, and Glenn C Turner. The Drosophila mushroom body: from architecture to algorithm in a learning circuit. Annual Review of Neuroscience, 43:465–484, 2020.

[5] Toshihide Hige, Yoshinori Aso, Mehrab N Modi, Gerald M Rubin, and Glenn C Turner. Heterosynaptic plasticity underlies aversive olfactory learning in Drosophila. Neuron, 88(5): 985–998, 2015.

[6] Kyle S Honegger, Robert AA Campbell, and Glenn C Turner. Cellular-resolution population imaging reveals robust sparse coding in the Drosophila mushroom body. Journal of Neuroscience, 31(33):11772–11785, 2011.

[7] Claire Eschbach, Akira Fushiki, Michael Winding, Casey M Schneider-Mizell, Mei Shao, Rebecca Arruda, Katharina Eichler, Javier Valdes-Aleman, Tomoko Ohyama, Andreas S Thum, et al. Multilevel feedback architecture for adaptive regulation of learning in the insect brain. bioRxiv, page 649731, 2019.

[8] Scott Waddell. Reinforcement signalling in Drosophila; dopamine does it all after all. Current Opinion in Neurobiology, 23(3):324–329, 2013.

[9] Yoshinori Aso, Daisuke Hattori, Yang Yu, Rebecca M Johnston, Nirmala A Iyer, Teri-TB Ngo, Heather Dionne, LF Abbott, Richard Axel, Hiromu Tanimoto, et al. The neuronal architecture of the mushroom body provides a logic for associative learning. Elife, 3:e04577, 2014.

[10] Trevor Hastie, Robert Tibshirani, and Jerome H Friedman. The Elements of Statistical Learning: Data Mining, Inference, and Prediction, volume 2. Springer, 2009.

[11] Donald Michie, David J Spiegelhalter, and Charles C Taylor, editors. Machine Learning, Neural and Statistical Classification. Ellis Horwood, 1994.

[12] Chanchal Chatterjee and Vwani P Roychowdhury. On self-organizing algorithms and networks for class-separability features. IEEE Transactions on Neural Networks, 8(3):663–678, 1997.

[13] Güleser Kalayci Demir and Kemal Ozmehmet. Online local learning algorithms for linear discriminant analysis. Pattern Recognition Letters, 26(4):421–431, 2005.

[14] Youness Aliyari Ghassabeh, Frank Rudzicz, and Hamid Abrishami Moghaddam. Fast incremental LDA feature extraction. Pattern Recognition, 48(6):1999–2012, 2015.

[15] Francis R Bach and Michael I Jordan. A probabilistic interpretation of canonical correlation analysis. Technical Report, 2005.

[16] Cengiz Pehlevan, Xinyuan Zhao, Anirvan M Sengupta, and Dmitri Chklovskii. Neurons as canonical correlation analyzers. Frontiers in Computational Neuroscience, 14:55, 2020.

[17] Siavash Golkar, David Lipshutz, Yanis Bahroun, Anirvan Sengupta, and Dmitri Chklovskii. A simple normative network approximates local non-hebbian learning in the cortex. Advances in Neural Information Processing systems, 33:7283–7295, 2020.

[18] David Lipshutz, Yanis Bahroun, Siavash Golkar, Anirvan M Sengupta, and Dmitri B Chklovskii. A biologically plausible neural network for multichannel canonical correlation analysis. Neural Computation, 33(9):2309–2352, 2021.

[19] Ramón Huerta and Thomas Nowotny. Fast and robust learning by reinforcement signals: explorations in the insect brain. Neural Computation, 21(8):2123–2151, 2009.

[20] Maxim Bazhenov, Ramón Huerta, and Brian H Smith. A computational framework for understanding decision making through integration of basic learning rules. Journal of Neuroscience, 33(13):5686–5697, 2013.

[21] James EM Bennett, Andrew Philippides, and Thomas Nowotny. Learning with reinforcement prediction errors in a model of the Drosophila mushroom body. Nature Communications, 12 (1):1–14, 2021.

[22] Robert AA Campbell, Kyle S Honegger, Hongtao Qin, Wanhe Li, Ebru Demir, and Glenn C Turner. Imaging a population code for odor identity in the Drosophila mushroom body. Journal of Neuroscience, 33(25):10568–10581, 2013.

[23] Ramón Huerta, Thomas Nowotny, Marta García-Sanchez, H. D. I. Abarbanel, and M. I. Rabinovich. Learning classification in the olfactory system of insects. Neural Computation, 16 (8):1601–1640, 2004.

[24] Darren Smith, Jan Wessnitzer, and Barbara Webb. A model of associative learning in the mushroom body. Biological Cybernetics, 99(2):89–103, 2008.

[25] Fei Peng and Lars Chittka. A simple computational model of the bee mushroom body can explain seemingly complex forms of olfactory learning and memory. Current Biology, 27(2): 224–230, 2017.

[26] Feng Li, Jack W Lindsey, Elizabeth C Marin, Nils Otto, Marisa Dreher, Georgia Dempsey, Ildiko Stark, Alexander S Bates, Markus William Pleijzier, Philipp Schlegel, et al. The connectome of the adult drosophila mushroom body provides insights into function. Elife, 9:e62576, 2020.

